# Machine learning methods trained on simple models can predict critical transitions in complex natural systems

**DOI:** 10.1101/2021.03.15.435556

**Authors:** Smita Deb, Sahil Sidheekh, Christopher F. Clements, Narayanan C. Krishnan, Partha S. Dutta

**Affiliations:** Department of Mathematics, Indian Institute of Technology Ropar, Rupnagar, Punjab 140 001 India; Department of Computer Science and Engineering, Indian Institute of Technology Ropar, Rupnagar, Punjab 140 001 India; School of Biological Sciences, University of Bristol, Bristol BS8 1TQ, UK

**Keywords:** alternative stable states, critical transitions, deep learning, early warning signals, machine learning

## Abstract

1. Sudden transitions from one stable state to a contrasting state occur in complex systems ranging from the collapse of ecological populations to climatic change, with much recent work seeking to develop methods to predict these unexpected transitions from signals in time series data. However, previously developed methods vary widely in their reliability, and fail to classify whether an approaching collapse might be catastrophic (and hard to reverse) or non-catastrophic (easier to reverse) with significant implications for how such systems are managed.

2. Here we develop a novel detection method, using simulated outcomes from a range of simple mathematical models with varying nonlinearity to train a deep neural network to detect critical transitions - the Early Warning Signal Network (EWSNet).

3. We demonstrate that this neural network (EWSNet), trained on simulated data with minimal assumptions about the underlying structure of the system, can predict with high reliability observed real-world transitions in ecological and climatological data. Importantly, our model appears to capture latent properties in time series missed by previous warning signals approaches, allowing us to not only detect if a transition is approaching but critically whether the collapse will be catastrophic or non-catastrophic.

4. The EWSNet can flag a critical transition with unprecedented accuracy, overcoming some of the major limitations of traditional methods based on phenomena such as Critical Slowing Down. These novel properties mean EWSNet has the potential to serve as a universal indicator of transitions across a broad spectrum of complex systems, without requiring information on the structure of the system being monitored. Our work highlights the practicality of deep learning for addressing further questions pertaining to ecosystem collapse and have much broader management implications.

## 1 Introduction

Transition from one steady state to another occur in many complex systems, such as financial markets (May et al 2008), human societies (Scheffer 2009, Downey et al 2016, Scheffer 2016), climate systems (Dakos et al 2008, Lenton 2011, Lenton et al 2019), systems biology (Sarkar et al 2019), and ecosystems (Scheffer and Carpenter 2003, Veraart et al 2012, Rindi et al 2017). Such transitions can be abrupt and irreversible (i.e., catastrophic), or smooth and reversible (i.e., non-catastrophic), and can occur due to gradual external forcing or random fluctuations in the system. In such scenarios, on crossing a threshold (known as a tipping or bifurcation point), structural changes occur in the underlying system. This is often termed a critical transition, prior to which the system’s return to an equilibrium slows down - a phenomenon known as critical slowing down (CSD) (Scheffer 2009). The phenomenon of CSD is related to the fact that the real part of the dominant eigenvalue of the system goes to zero at the bifurcation point (Wissel 1984, Scheffer et al 2009). In all such cases, where the dominant eigenvalue approaches zero close to the tipping point irrespective of catastrophic or non-catastrophic transitions, the phenomenon of CSD persists, and there exist statistical indicators that forewarn the vicinity of a tipping point (Kéfi et al 2013). Understanding the causes of sudden transitions and forecasting them using statistical indicators have recently emerged as an important area of research due to the management implications of preventing catastrophes in natural systems (Scheffer et al 2009, 2012, Dakos et al 2012a).

The traditional approach of forecasting a critical transition relies on summary statistics such as variance, autocorrelation, and skewness showing an increasing trend before a transition. Although model simulations and experiments have demonstrated the usefulness of such indicators, the robustness of generic early warning signals (EWSs) in forecasting critical transitions remains debatable (Boettiger and Hastings 2012b,a, Clements et al 2015, Ditlevsen and Johnsen 2010). The uncertainties in generic EWSs can be attributed to factors including imperfect data sampling, lack of quantitative and objective measures, short time series, and sensitivity to bandwidth and window sizes. However, despite these challenges, significant work has sought to increase the statistical power of the EWSs in the hope they can be used as predictive management tools. This includes recent work showing that incorporating additional measures of the health of a system (such as the mean body size of biological populations) can increase the efficiency in predicting a tipping point (Clements and Ozgul 2016, Clements et al 2017, Drake and Griffen 2010). Whilst such approaches are promising, they require more data than traditional generic EWSs (two or more simultaneous time series), limiting their applicability to many systems (Clements and Ozgul 2016). Consequently, there remains a critical need to develop a robust toolkit to identify tipping points using widely available time series data. Were this to be achieved, it could have significant implications for the management of a host of systems, from financial markets to species at risk of extinction (Sarkar et al 2019, Scheffer and Carpenter 2003, Veraart et al 2012).

One potentially powerful tool to achieve this is machine learning (ML). ML models are able to automatically capture statistical characteristics by identifying and learning hidden patterns in data (Pathak et al 2018), making them ideally suited to detecting warning signals. Indeed, ML has been used to classify phases of matter, study phase behavior, detect phase transitions, and predict chaotic dynamics (Scandolo 2019, Van Nieuwenburg et al 2017, Canabarro et al 2019, Zhao and Fu 2019, Pathak et al 2018), whilst supervised learning algorithms such as artificial neural networks have been used to study second-order phase transitions, especially the Ising model (Morningstar and Melko 2017, Cossu et al 2019, Ni et al 2019, Giannetti et al 2019). However, thus far machine learning tools have not been used to classify the most common transitions seen in ecological, financial, and climatic systems - catastrophic (i.e., first-order or discontinuous) and non-catastrophic (i.e., second-order or continuous) transitions (Martín et al 2015).

In this study, we build a deep learning model, the Early Warning Signal Network (EWSNet), and train it on the raw time series data simulated from nine different dynamical models, including biological, ecological, and climate models displaying a range of sudden, smooth, and no transitions (see *SI Appendix*, Section S1 and Table S1). We show that EWSNet provides ro-bust identification of approaching tipping points in simulated data, and that it outperforms four classical ML models (logistic regression, support vector machine (SVM), random forest, and multilayer perceptron (MLP) (Pichler et al 2020, Olden et al 2008, Bishop 2006)) which are trained to classify time series based on trends in their statistical properties such as autocorrelation - the basis for generic early warning signals (Dakos et al 2012a). Furthermore, we then show that, even though EWSNet is trained on simulated time series data, it can reliably predict approaching transitions in seventeen real-world and experimental data sets (Clements and Ozgul 2016, Clements et al 2017, Dakos et al 2008, Chen et al 2018). Our results suggest that EWSNet identifies information hidden implicitly in time series that is missed by generic EWSs, and because of this EWSNet can reliably predict the future state of a range of complex systems even when time series are imperfectly sampled (Clements et al 2015). The approach of the EWSNet as an early warning indicator makes few assumptions about the underlying state of the system, requires no data pre-processing, and is invariant to sequence length, and thus offers a novel and powerful predictive management tool.

## 2 MATERIALS AND METHODS

### 2.1 Generation of simulated training data

We have generated stochastic time series from nine different models ranging from ecological to paleoclimatic systems that cover a wide range of nonlinearities (see *SI Appendix*, Section S1). The considered models are of the form:

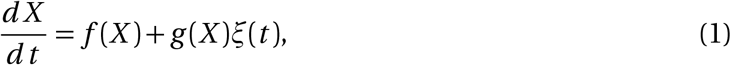

where *X* is the state variable, *f* (*X*) is the deterministic skeleton of the model, *g* (*X*) is an arbitrary function, and *ξ*(*t*) is a random variable depicting colored noise. The effect of colored noise was incorporated in the deterministic skeleton by the equation:

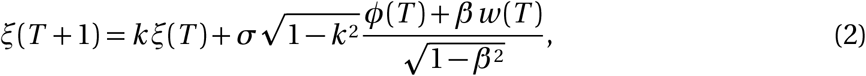

where 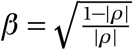, is the species response correlation, *T* represents the time points (1,....,400), *k* is the autocorrelation coefficient, and *σ* is the noise intensity. *ϕ*(*T*) and *w* (*T*) are normal random components, where *w* (*T*) differs across species unlike *ϕ*(*T*) (Yang et al 2019, Ruokolainen 1980). The stochastic models were simulated using the Euler-Maruyama method with *k* ∈ [−0.8, 0.8]. We majorly trained our deep neural network using a large number of simulated time series data perturbed with white noise (i.e., *k* = 0). For testing our model potency in an-ticipating transitions in time series for systems perturbed with multiplicative noise, we train our model with an additional number of time series which have been perturbed with coloured noise. This is done in order to let the EWSNet be familiar with the fast changing dynamics and short scale fluctuations (Ditlevsen and Johnsen 2010) that occur due to the presence of multiplicative noise, and also to test the skills of EWSNet in testing datasets which may not fall in the known regimes of training data.

### 2.2 Deep learning model: EWSNet structure

The EWSNet (see Fig. 1) can be viewed as a large composition of complex nonlinear functions that learn hierarchical representations of the data. The input to the EWSNet is a univariate time series signal. The EWSNet comprises of FCN, and LSTM blocks, followed by fully connected layers (Karim et al 2018). The FCN consists of three stacked convolutional blocks, each composed of convolution (Goodfellow et al 2016), batch normalization (BN) (Ioffe and Szegedy 2015), and rectified linear unit (ReLU) (Nair and Hinton 2010) activation layers. The convolution operation is performed using a filter **W** ∈ ℛ^1×*k*^ over an input tensor **X** ∈ ℛ^1×*T*^, where *T* is the length of the time series. These filters are learnable and often characterize various local patterns present in the input tensor. The convolution operation is followed by batch normalization to remove the covariate shift in the output across different training batches. The ReLU activation function is applied to the batch normalized output. The resulting output at the end of one convolution block can be represented as:

**Figure 1:**
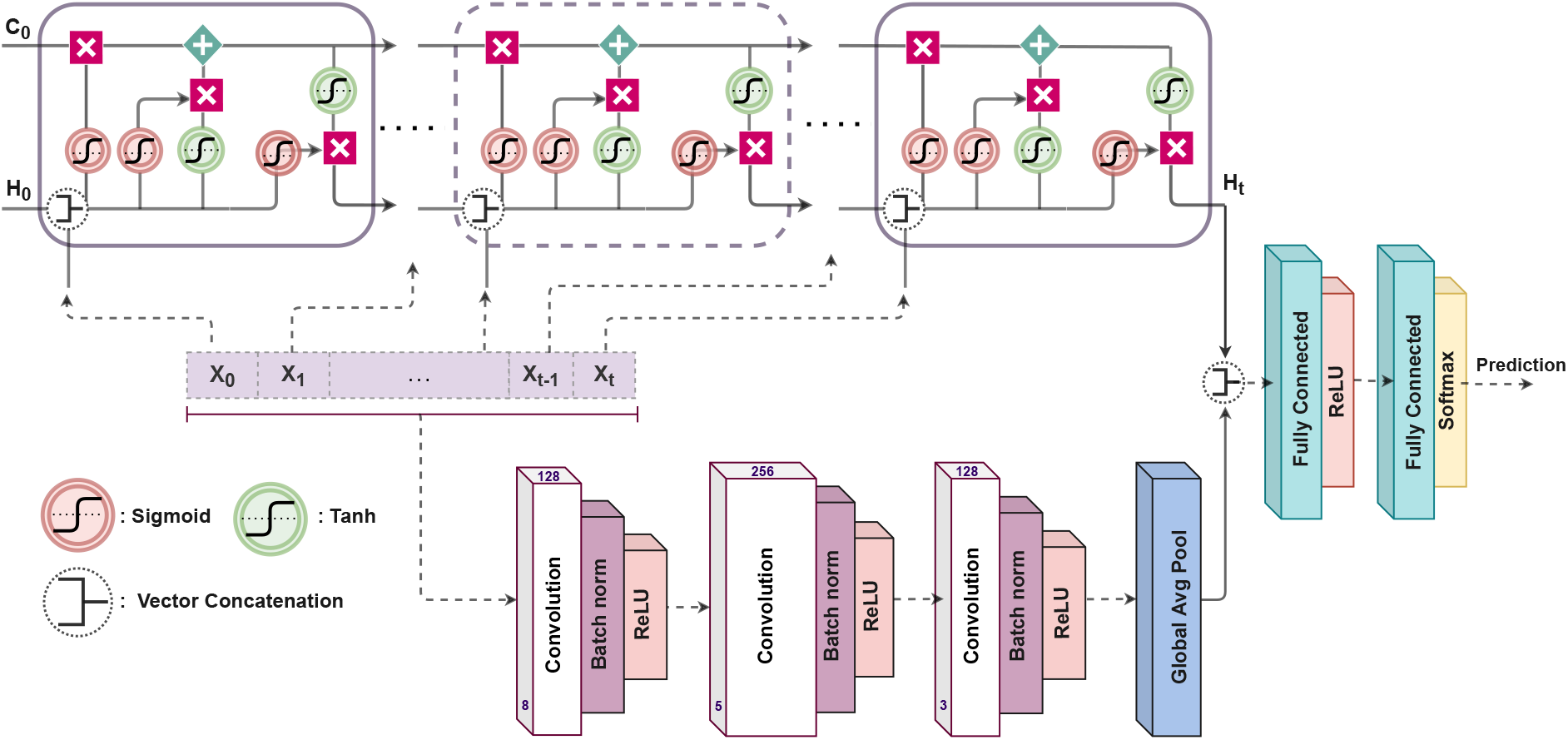
Schematic representation of the EWSNet: The EWSNet consists of three convolution blocks and an LSTM block. The fully convoluted network and the LSTM block process the input sequence independently. The concatenated output of the two blocks is passed through two fully connected layers to obtain the final prediction. X_*t*_ represents the input at time step *t*. C_0_ and H_0_ represent the initial cell and hidden states of the LSTM block, respectively. The cell state allows the passage of stored information, and the hidden state acts as the working memory to the LSTM block of the EWSNet. The Global Average Pooling layer at the end of the convolutional block makes the EWSNet invariant to sequence length.

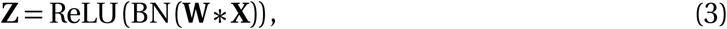

where * represents the convolution operation. Each of the three convolution blocks of EWSNet processes the output of the previous block in a similar fashion. The output of the third convolutional block, containing *D* filters, is a set of *D* vectors, each of length *T*. To make EWSNet invariant to the sequence length *T*, we apply a global average pooling operation (over *T*) to obtain a *D* dimensional vector.

An **LSTM** (Hochreiter and Schmidhuber 1997) is a recurrent neural network that integrates a gradient superhighway in the form of a cell state **C**_*t*_, in addition to the hidden state **H**_*t*_ at time *t*. The LSTM model has gates which allow addition and removal of information to and from the cell state. The forget gate (**f**_*t*_), parameterized by **w**_*f*_, decides the information to be deleted from the cell state **C**_*t*_ on obtaining the input **X**_*t*_ at time *t* and can be defined as follows:

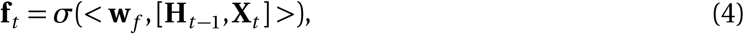

where *σ* is the sigmoid activation function and <. > is the dot product operation. The input gate (**i**_*t*_), parameterized by **w**_*i*_, determines the information that should be added to the cell state **C**_*t*_ and is defined as:

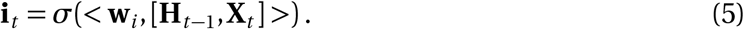

The cell state **C**_*t*_ is obtained by using both **f**_*t*_ and **i**_*t*_ in the following manner:

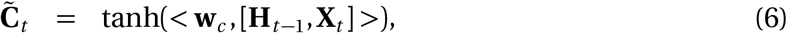

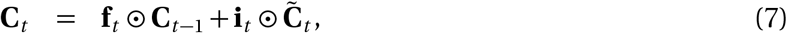

where **w**_*c*_ parameterizes the intermediate cell state and 0 is the Hadamard product. The hidden state **H**_*t*_ and output gate **o**_*t*_ (parameterized by **w**_*o*_) of the LSTM are defined as:

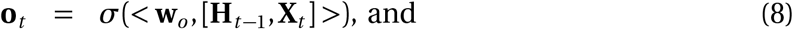

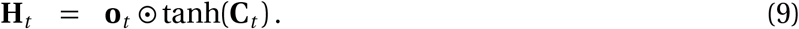

The EWSNet concatenates the hidden state of the LSTM after observing the last input with the output of the FCN. This concatenated feature representation of the input time series is used to obtain the final output after passing through two fully connected layers. A fully connected layer consists of multiple nodes, each of which is connected to every output of the previous layer. Thus, each node (parameterized by **w**) takes a vector, **x**, as input and outputs a scalar that is a nonlinear transformation of the weighted sum of the inputs in the following manner:

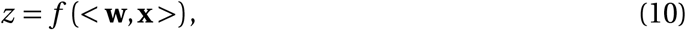

where *f* (·) is a non-linear transformation. The first fully connected layer of the EWSNet uses ReLu as the activation function, while the second layer uses Softmax activation resulting in a probability distribution over the class labels. The parameters of EWSNet at each layer are learned using the backpropagation algorithm to minimize the overall cross-entropy loss be-tween the predictions and the ground truth. The EWSNet comprises 3 convolution layers with 128, 256, and 128 filters of size 3, 5 and 3 respectively, and an LSTM block with a hidden state of dimension 128. These choices for the hyper-parameters were obtained through fine-tuning (see *SI Appendix*, Section S1, Fig. S2). We used an ADAM (Kingma and Ba 2015) optimizer with the learning rate 5 × 10^−5^ and batch size 512 to train the EWSNet.

### 2.3 Machine learning models trained on generic EWSs

We use classical ML models, namely Logistic Regression, Random Forest, SVM, MLP, to classify time series based on the extracted EWSs passed as input to these models. Generic EWSs are calculated using time series for a combination of bandwidths ({20, 30, 40}) and window sizes ({40, 50, 60}). This is done with the idea that the EWSs for a particular window size and bandwidth capture certain characteristic features that aid in classifying the time series and assigning the appropriate label (see *SI Appendix*, Section S2). Accordingly, if the individual EWSs for the combinations are concatenated and passed as input to these models, they act as an additional filter to the generic EWSs and are supposed to improve them. A brief description of the model along with the tuned hyperparameters for the respective models are discussed in *SI Appendix*, Section S2, Table S2.

## 3 Results

We have engineered a deep learning model - EWSNet to forewarn an upcoming shift on simulated and real-world time series data. The EWSNet (see Fig. 1) consists of two blocks; the LSTM block that captures the latent temporal properties and the fully convolutional block that views the time series as spatial data. The hyper-parameters of the model (such as the number of convolution blocks, LSTM hidden units, learning rate, etc.) were carefully fine-tuned using the training and the validation sets (see *Materials and Methods*). Two experimental regimes derived from nine different models pertaining to biological, ecological, and paleoclimatic dynamics (Scheffer et al 2012, Sharma and Dutta 2017, Dutta et al 2018, Boerlijst et al 2013, Dakos et al 2008) (see *SI Appendix*, Table S1) perturbed with white noise (Dataset-W) and colored noise (Dataset-C) are used to train and test the performance of EWSNet. The time series exhibit catastrophic and non-catastrophic transitions with CSD (spanning over four different bifurcations; viz, saddle-node (fold), transcritical, pitchfork, and supercritical Hopf bifurcations) and no transitions. We have also included in the study time series with coloured noise where the generic EWSs typically show weak trends (Seekell et al 2011, Rudnick and Davis 2003, Dutta et al 2018). To estimate the robustness of the models and to rule out results due to chance, we report the performance of the models averaged over 25 trials.

### 3.1 Detecting and characterizing transitions using EWSNet

Two different EWSNet models were each trained independently on Dataset-W and Dataset-C, respectively using 80% of the generated dataset. The remaining 20% was used for testing the performance of the trained model. Further, the test set of Dataset-C contained time series with highly correlated noise, while the training set contained time series with weakly correlated noise. The training and validation accuracies for both the models as a function of the number of training epochs averaged over 25 different trials are presented in Fig. 2. We observe the convergence of the model after 25 epochs. We also noticed transients in the training epochs that are symptomatic of training deep neural network models. The mean test accuracy of the EWSNet models for Dataset-W is 99.46% and for Dataset-C is 95.93%. As can be othe accuracies of the generibcserved from Fig. 3, the EWSNet models show high accuracy for all the three labels (catastrophic (C.T.), non-catastrophic/smooth (S.T.), and no transition (N.T.)) on both the datasets. While non-transition series are always correctly classified by both the models, there exists a small amount of confusion for the other two classes. Moreover, the efficacy of EWSNet for time series of varying lengths and at varying distances from the tipping point (if present) are discussed in the supplementary material (see *SI Appendix*, Section S1, Fig. S1). The generic EWSs are sensitive to the length of pre-transition time series used, whereas the EWSNet models are marginally affected by reduced time series length.

**Figure 2:**
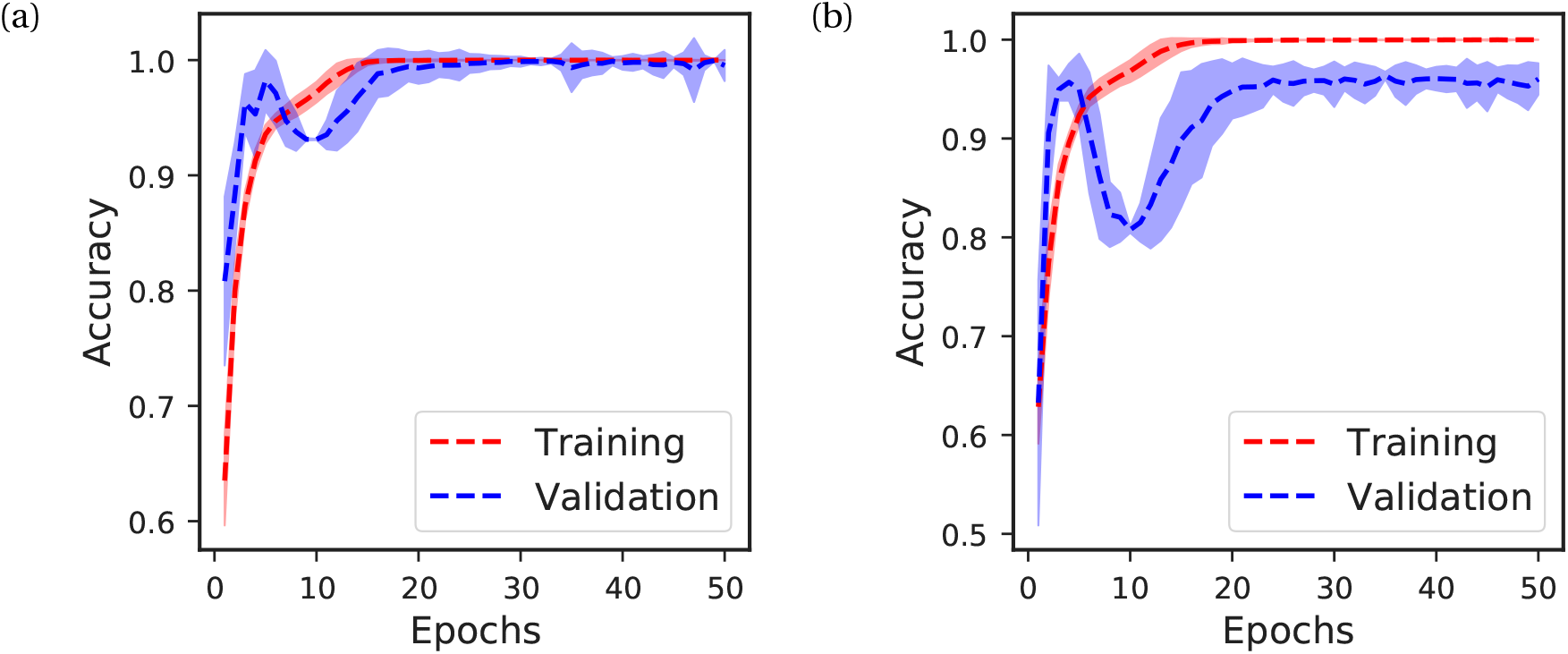
Mean accuracy of EWSNet: for (a) Dataset-W, and (b) Dataset-C. The mean training and validation accuracies are computed after every epoch and are averaged over 25 trials. The shaded regions represent the 95% confidence interval. The accuracies saturate after around 25 epochs indicating the models’ convergence.

**Figure 3:**
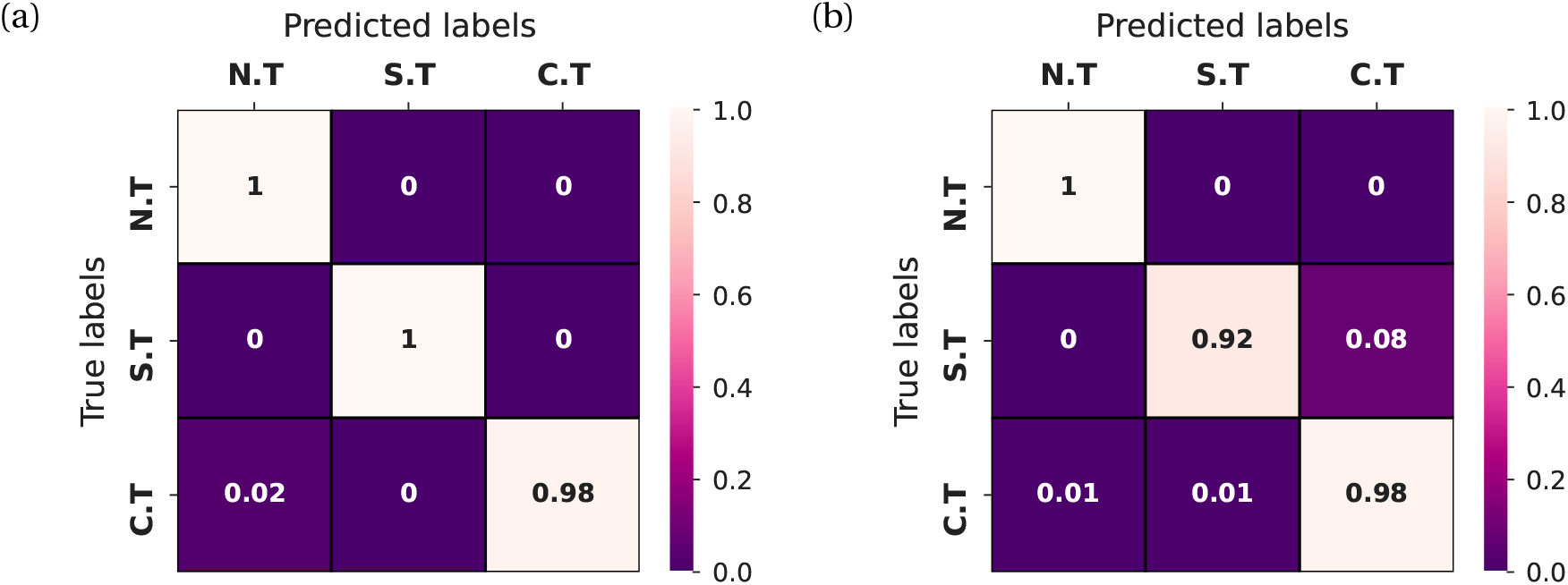
Error analysis using the confusion matrix: for (a) Dataset-W, and (b) Dataset-C. The EWSNet always classifies a no transition time series accurately. It rarely makes an error in classifying catastrophic and non-catastrophic transitions for Dataset-W. For Dataset-C, 1% of catastrophic transitions are labeled as non-catastrophic, and 8% of non-catastrophic transitions are being misclassified as catastrophic. Here, C.T., S.T., and N.T. stand for a catastrophic, non-catastrophic, and no transition, respectively.

Detection of EWSs using classical trends in statistics such as autocorrelation can be highly sensitive to non-uniform sampling of data (Boettiger and Hastings 2012b, Clements et al 2015), with sporadic sampling increasing the probability of misidentifying catastrophic and non-catastrophic transitions, or failing to detect a transition at all. We investigated the robustness of the trained EWSNet models towards imperfectly sampled time series by re-sampling the time series (see *SI Appendix*, Section S1). As expected, this imperfect sampling reduced the efficiency of generic EWSs. However, EWSNet models continued to provide robust predic-tions of approaching transitions; even when re-sampling retained only 40% of the original simulated data, the mean accuracy remained above 80% (see Fig. 4).

**Figure 4:**
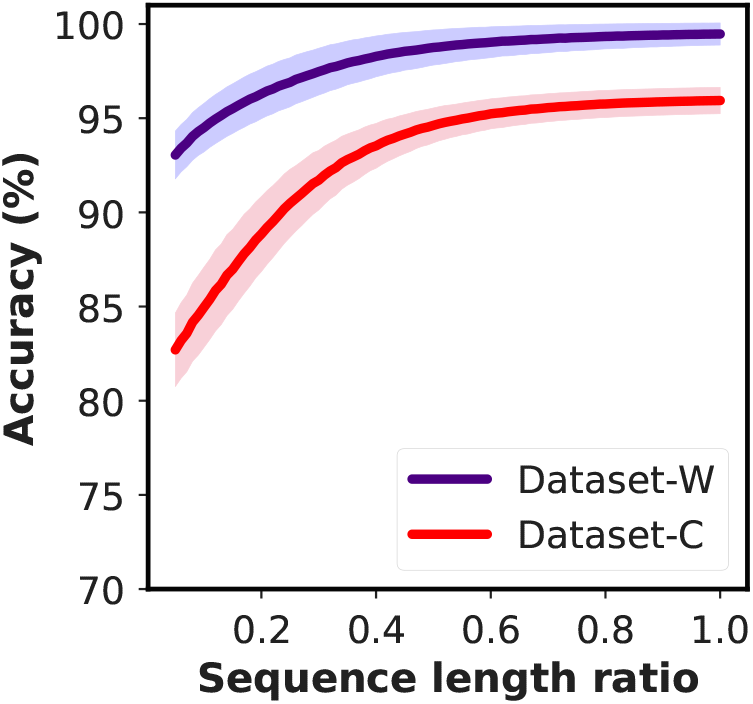
Robustness of EWSNet on imperfectly sampled data: EWSNet is robust to observational error. The model performance improves with an increase in the sequence length; the model performs well even for moderate infrequent sampling for both Dataset-W and Dataset-C. The infrequent temporal sampling is achieved by randomly selecting a subset of the original time series (increments of 0.01 to 1 of the actual length).

### 3.2 Performance of ML models trained using generic EWSs

Whilst EWSNet is trained on the raw simulated time series data, it is also possible to train ML models on the trends in generic early warning signals calculated from these simulations and thus use ML to attempt to detect similar trends in statistics such as autocorrelation in the validation data sets. To compare the efficacy of this approach to the EWSNet approach, we used four standard ML models (Giannetti et al 2019, Pichler et al 2020, Olden et al 2008) (logistic regression (LR), support vector machine (SVM), Multilayer perceptron (MLP), and random forests (RF)) to characterize trends in generic EWSs in simulated time series. The input to the ML model is the concatenation of the generic EWSs captured for different combinations of bandwidths and window-sizes (for details see *SI Appendix*, Section S2). The results presented in Fig. 5 show that AR-1 and SD are the top two performing generic EWSs across both the datasets and across all the ML models, in accordance with prior literature (Dakos et al 2012b). Thus, we further trained ML models using a combination of AR-1 and SD which resulted in performance greater than the models trained on the individual EWSs (see Fig. 5) and other pair-wise combinations of EWSs (see *SI Appendix*, Section S2, Figs. S3-S5).

**Figure 5:**
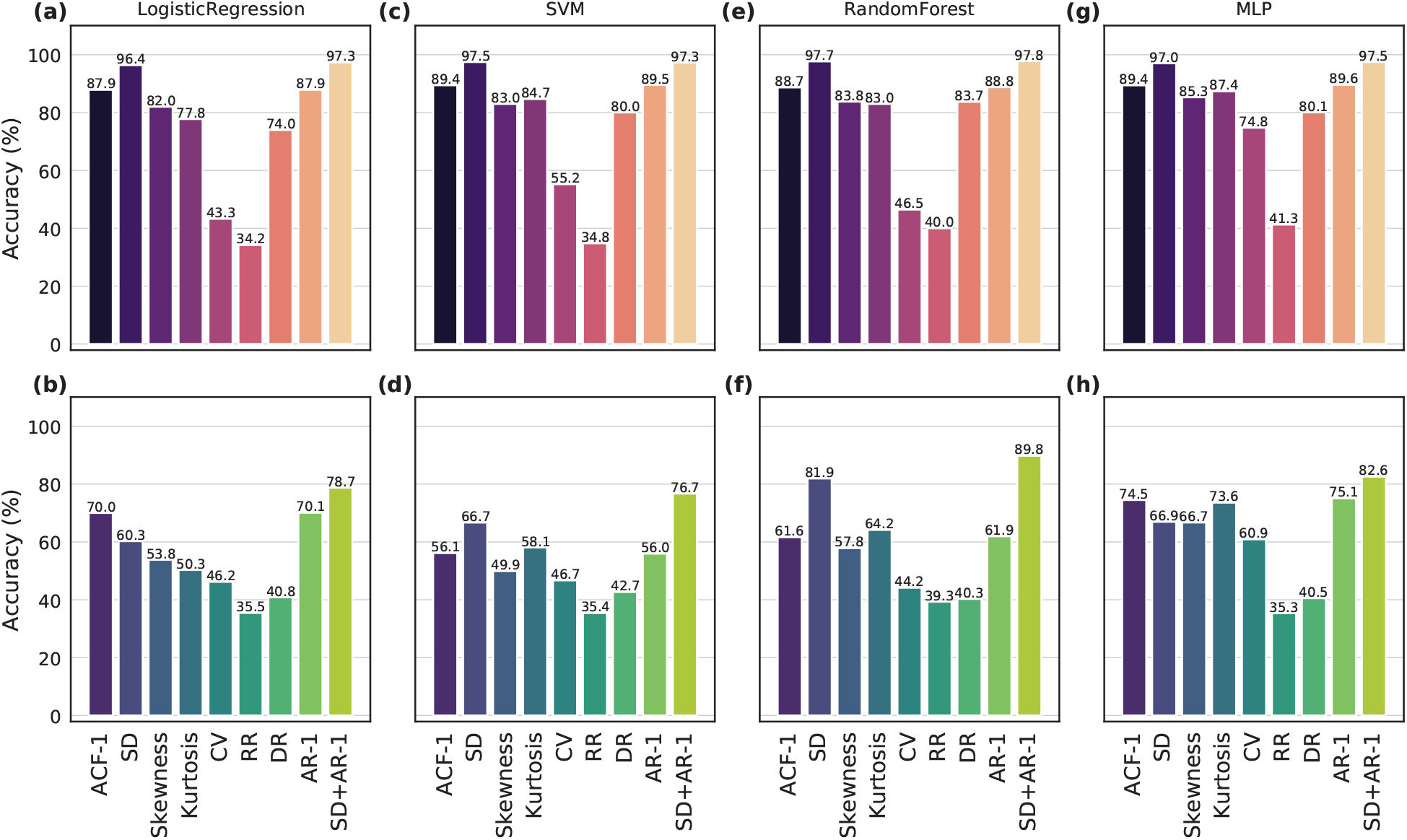
Performance of generic EWSs using (a,b) Logistic Regression, (c,d) Support Vector Machine (SVM), (e,f) Random Forest, and (g,h) Multilayer Perceptron (MLP): The ML models are trained on EWSs extracted using various combinations of rolling window sizes and bandwidths. The results are presented as bar plots corresponding to each EWS indicator for the Dataset-W (upper panel) and Dataset-C (lower panel). Among the best performing indicators are SD, AR1, and SD+AR1. The SD+AR1 model improves the performance over the individual SD and AR1 models.

Table 1 compares the accuracies of the generic EWSs based ML models against the EWSNet for both the Dataset-W and Dataset-C. EWSNet performs significantly better in distinguishing transitions and classifying time series for both the datasets: the probability for the *t*-statistics of obtaining a mean accuracy as that of EWSNet is negligible (*p* < 1 × 10^−5^). Interestingly, irrespective of the ML model chosen - tipping points are more accurately identified in Dataset-W than Dataset-C. This can be explained by the short-scale fluctuations introduced by colored noise, in Dataset-C, which can dampen the trend in the time-series.

**Table 1:** Comparison of mean accuracy for various models: On comparing the mean accuracy, EWSNet appears to be the best performing model in classifying a critical transitional time series consistently for both the Dataset-W and Dataset-C. On passing the EWSs as input to the other ML models, the results are comparably close to the EWSNet for Dataset-W, but the accuracy declines for Dataset-C. The mean accuracy for the four classical ML models is comparable. The random forest is the best among them, with standard deviation and a combination of standard deviation and autocorrelation at lag-1 as features. The numeric value in ± denotes 95% confidence interval.

| ML methods | EWSs | Dataset-W | Dataset-C |
| --- | --- | --- | --- |
| Support Vector Machine | SD | $97.47 \pm 0.09$ | $66.69 \pm 0.00$ |
| | AR1 | $89.55 \pm 0.17$ | $55.95 \pm 0.00$ |
| | SD + AR1 | $97.27 \pm 0.14$ | $76.70 \pm 0.00$ |
| Logistic Regression | SD | $96.40 \pm 0.14$ | $60.27 \pm 0.01$ |
| | AR1 | $87.88 \pm 0.22$ | $70.08 \pm 0.02$ |
| | SD + AR1 | $97.35 \pm 0.11$ | $78.74 \pm 0.02$ |
| Random Forest | SD | $97.69 \pm 0.12$ | $81.93 \pm 0.22$ |
| | AR1 | $88.76 \pm 0.20$ | $61.94 \pm 0.30$ |
| | SD + AR1 | $97.77 \pm 0.16$ | $89.79 \pm 0.82$ |
| Multilayer Perceptron | SD | $97.02 \pm 0.69$ | $66.94 \pm 1.55$ |
| | AR1 | $89.58 \pm 0.45$ | $75.10 \pm 0.82$ |
| | SD + AR1 | $97.47 \pm 0.16$ | $82.56 \pm 2.10$ |
| EWSNet | | $99.46 \pm 0.01$ | $95.93 \pm 0.02$ |

Populations experiencing red noise are relatively more threatened and found to encounter long periods of survival conditions (Vasseur and Yodzis 2004, Yang et al 2019) and their transitions need to be anticipated. Therefore, we compared the EWSNet against the ML models on Dataset-C (the test set contains time series with high correlated noise). The metrics Accuracy, Area Under the Curve (AUC), True Positive Rate (TPR), Precision, and F Score for each of the models are presented in Fig. 6, again EWSNet outperforms other methods for these red noise time series.

**Figure 6:**
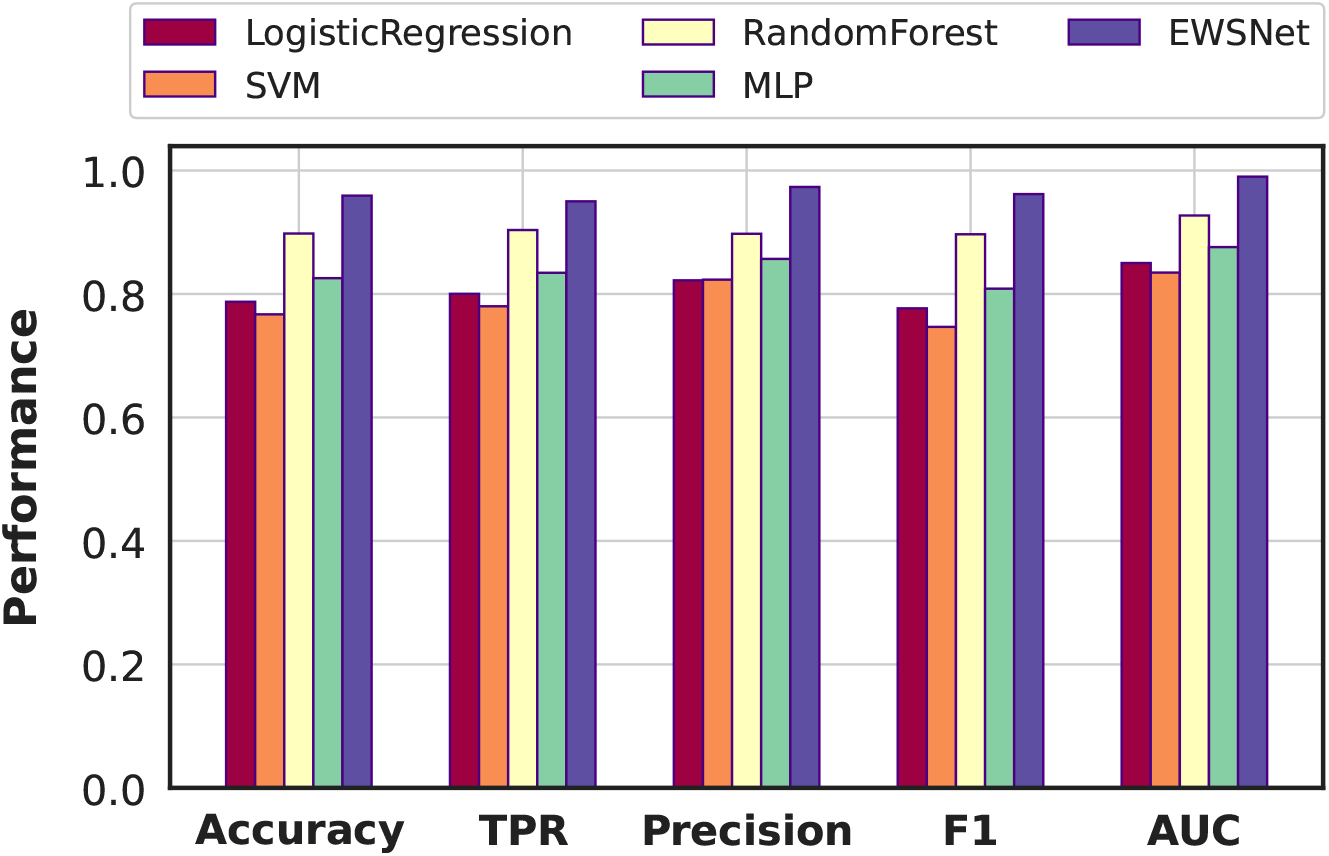
Predictive performance of different models: Comparison of Logistic Regression, SVM, Random Forest, MLP, and EWSNet across five metrics, namely Accuracy, TPR, Precision, AUC, and F1 Score. EWSNet results in the best performance uniformly across all the metrics.

### 3.3 Applicability and robustness of EWSNet to real world and experimental data

The noisy nature of non-simulated data means that the effectiveness of EWSs is often variable (Burthe et al 2016). To test the reliability of EWSNet when predicting real world data, we have considered seventeen ecological (Chen et al 2018, Clements and Ozgul 2016, Clements et al 2017) and paleoclimatic data (Dakos et al 2008) (see *SI Appendix*, Section S3, Figs. S6-S7). We used the EWSNet trained with simulated data and tested it on the above real time series. Applying classical ML models on real time series that vary in length, some being inexplicably short, is challenging and requires additional effort to make them of equal length. The additional steps will either pad, interpolate, or truncate the sequences leading to information loss in the trend and erroneous results. In contrast, EWSNet can classify time series of varying lengths due to the global average pooling post convolution layers, making the model dynamic. We present the prediction probabilities for the EWSNet, and Kendall’s-*τ* correlation coefficient for the generic EWS (AR-1) (see *SI Appendix*, Table S3). The EWSNet classifies and distinguishes transitions in real time series with a mean accuracy of 76.47 ± 0.03 over 25 trials with high predictive probability (see Fig. 7). This is the first time the results have been computed on real-world time series objectively using deep learning models trained only on simulated time series.

**Figure 7:**
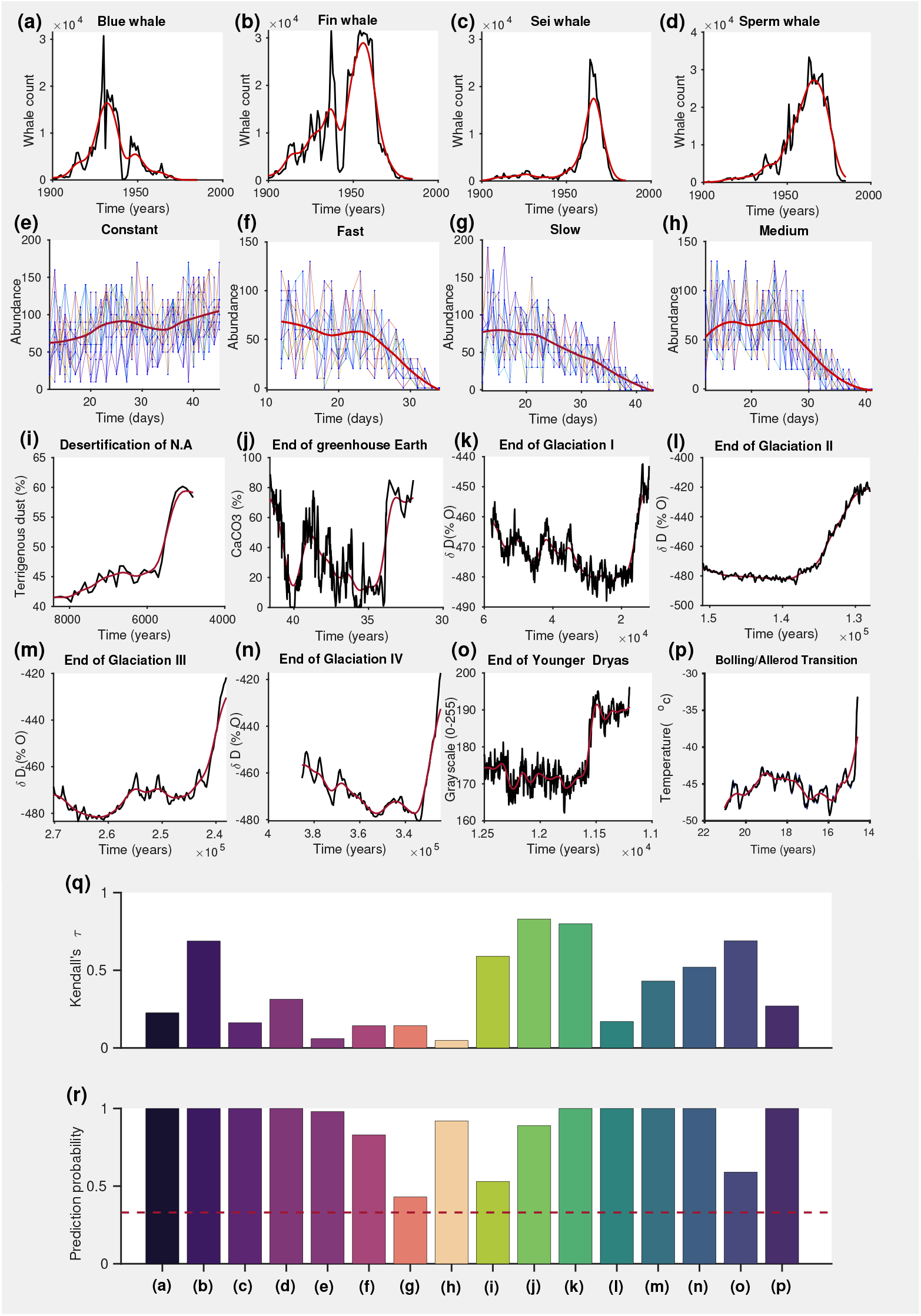
Performance on real world and experimental time series: Time series data of: (a)-(d) different whale counts, (e)-(h) abundance of *Didinium nasutum* populations that were exposed to four different experimental treatments, and (i)-(p) climate systems. In (a)-(p), the red curve is the time series trend. Histograms showing performance of (q) AR-1 using Kendall’s-*τ*, and (r) the EWSNet using prediction probability, respectively, for the time series data ((a)-(p)). In (r), the red dashed line denotes prediction probability due to chance. The results for dryland ecosystem are presented in *SI Appendix*, Table S3.

## Discussion

Previously developed EWSs (Scheffer et al 2009, Drake and Griffen 2010, Clements and Ozgul 2016) have had limited success in predicting approaching transitions, particularly in noisy real world data. Moreover, such methods cannot discern whether an approaching transition is catastrophic or not. Here we take an entirely new approach to predict transitions in complex systems by developing a deep learning model (EWSNet) that can forecast not only an approaching tipping point but also discern whether it is catastrophic or not. The key characteristic of our model is its non-dependence on a priori statistical features such as increasing autocorrelation in a time series. Rather, EWSNet unbiasedly learns latent features embedded in time series, which are indicative of approaching collapse. Using this approach EWSNet - when trained only on data generated from nine simple mathematical models - can yield high accuracy inferences of approach tipping points in noisy real-world and laboratory datasets.

The EWSNet has been trained using simulated time series data and results in a test accuracy of 99.46%±0.01 on the Dataset-W. The trained EWSNet achieves an approximate baseline accuracy of 90% (80%) when presented with only 5% of the time series data from Dataset-W (Dataset-C). The performance steadily improves with the availability of more information on the time series saturating at 99% (92%) for Dataset-W (Dataset-C) (Fig. 4). Thus the EWSNet reduces the need for high-frequency data while uncovering the complex dynamics to classify an impending transition. Adding a small number of time series with colored noise (correlation time < 0.2) during training results in EWSNet that achieves a mean test accuracy of 95.93% on a more challenging time series containing colored noise (0.2 < correlation time ≤ 0.8) (Table 1). Analyzing the false positives and negatives for each label, we perceive that the EWSNet never misclassifies a time series with no transition but infrequently confuses between catastrophic and non-catastrophic transitions (Fig. 3).

Next, we investigate the effectiveness of four ML models (both linear and non-linear) trained on trends in generic EWSs calculated from the simulated datasets. ML models trained using EWSs perform well for Dataset-W but suffer a significant drop in the performance on Dataset-C (see Fig. 6). Among the best-performing indicators are standard deviation, auto-correlation at lag-1, and a combination of these two, which is consistent with the findings in the literature (Dakos et al 2012b). The results are comparable to EWSNet for Dataset-W. However, EWSNet performs better than the generic EWSs based ML models on Dataset-C (see Table 1). The generic EWSs in a time series are insufficient to anticipate and classify the transitions. Therefore, we infer that the EWSNet captures other latent properties of the time series that the generic EWSs do not characterize. Further, classical approaches using generic EWSs require a careful selection of suitable bandwidth, and window-size (Lenton et al 2012, Boettiger and Hastings 2012b). However, the EWSNet does not have this limitation.

We also test the EWSNet, trained on simulated data, on real world time series containing both abrupt and smooth shifts. Based on the comparison of the predictive probabilities and the Kendall-*τ* values, we conclude that our model is efficient and more potent in forecasting a critical transition (see Fig. 7). The likelihood of the EWSNet predicting the true label is significantly greater than chance even for imperfectly sampled or shorter time series. However, the traditional methods based on generic EWSs can be susceptible to false positives for such time series (Dakos et al 2008).

Thus, we have engineered and demonstrated a deep neural network-EWSNet that results in better performance over methods utilizing generic EWSs and can be further developed to serve as a universal indicator of critical transitions. Considering transitions entailing colored noise, the usefulness of EWSNet increases manifold as it can learn the short-time scale fluctuations and classify a time series as a critical transitional or not. On the other hand, the generic EWSs do not exhibit a trend in cases when transitions occur due to chance fluctuations (Ditlevsen and Johnsen 2010). There remain a few open questions related to the detection of critical transitions in complex systems, such as machine learning models’ ability to detect bifurcations without CSD (non-smooth) and sharp transitions without bifurcations (Dutta et al 2018). Our work also shows promising future directions to develop machine learning models for classifying bifurcations and further predicting ‘when’ the critical transition will occur.

However, on a broader perspective, the EWSNet is a more generalized model with almost no presumptions opening up the possibility of real-time monitoring of many real-world systems such as global climate, lake ecosystem, cell dynamics in various organisms that generate distorted and noisy sequences of events for automatically predicting impending transitions with a negligible computational cost.

## Acknowledgments

S.D. acknowledges the Ministry of Education (MoE), Govt. of India for Prime Minister’s Research Fellowship (PMRF). P.S.D. acknowledges financial support from Science & Engineering Research Board (SERB), Govt. of India [Grant No.: CRG/2019/002402]. N.C.K acknowledges to resources and support received under Google TensorFlow Research Award.

## Authors’ contributions

N.C.K. and P.S.D. conceived the study. S.D. and S.S. performed the simulations and analyzed the results. C.F.C. performed the microcosm experiments. All authors discussed the results and wrote the manuscript.

## Supporting Information

Additional supporting information is available with this article.

