## SUPPLEMENTARY MATERIAL for "Machine learning methods trained on simple models can predict critical transitions in complex natural systems"

**Contents**

**S1. Data generation for the EWSNet** **2**

**S2. Data generation for classical ML models trained on generic EWSs** **7**

**S3. Real data description** **10**

### S1. Data generation for the EWSNet

To study the functionality of the EWSNet as an indicator to forewarn sudden transitions in complex systems, we generate time series by simulating nine well studied models that exhibit transitions via four different types of co-dimension one bifurcations, i.e., saddle-node (fold), transcritical, pitchfork, supercritical Hopf, and also no transition (see Table S1). We consider different nonlinear models that can undergo saddle-node bifurcation and have alternative stable states within a range of values of the control parameters, which is considered as a catastrophic transition (i.e., first-order or discontinuous transition). The other models exhibit transcritical, pitchfork, and supercritical Hopf bifurcations and several time series have been generated from them are categorized to belong to the class of non-catastrophic transitions (i.e., second-order or continuous transitions). We also consider the first model and vary the control parameter in a range such that there is no transition (neither catastrophic nor non-catastrophic) (Table S1 model [9]). The first model (Table S1 model [1]) is a paradigm of catastrophic transitions in an ecosystem (Ludwig et al 1978). It depicts the growth of a resource ($N$) and its consumption under the effect of immense grazing pressure, there occurs a catastrophic shift in the resource. The other two ecological models considered for catastrophic transitions are a model depicting overexploitation (Table S1 model [2]) and a juvenile-adult model (Table S1 model [3]) (Scheffer et al 2012, Boerlijst et al 2013). Next, we consider a bistable gene regulatory positive feedback loop model (Table S1 model [4]) depicting the synthesis of protein ($x$) in microorganisms that break down nutrients to release energy to the cell (Cheng et al 2008). Sudden transitions in protein concentration can be a cause of complex disease emergence. We also consider a complex climate model (Table S1 model [5]) to make the EWSNet acquainted with dynamics of abrupt climate shifts (Dakos et al 2008, Fraedrich 1978). Further, we pick three models (Table S1 models [6]-[8]) of non-catastrophic transitions. Particularly, model [8] may exhibit a non-catastrophic transition from a steady-state to an oscillatory state due to interspecific competition between consumer and resource. Here, we consider different models pertaining to each transition variant to make the EWSNet familiar with a range of nonlinearities. For each model, we choose different values of a parameter along with the control parameter (which is the driver of a transition) to bring in maximum possible variations across time series preserving the aforementioned bifurcations.

We solve the models [1]-[9] (see Table S1) using the Euler-Maruyama method with an integration step size $dt = 0.01$. In all the above models with their deterministic counterparts we consider multiplicative noise (see "Methods" in the main draft), and then simulate the corresponding stochastic systems over a range of stochasticity covering the noise color spectra with $\kappa \in [-0.8, 0.8]$, to obtain time series. We vary the noise intensity $\sigma$ in the range $(0.03, 0.07)$. The standard normal components in Eq. (1) (in the main draft), i.e., $\phi(t)$ and $\xi(t)$ add in a wide variety of stochasticity in the systems which when used to train the EWSNet can make the model familiar with features particular to different noise colors (Yang et al 2019).

For time series that are generated by perturbing the system with positive correlations, the tipping point precedes the deterministic bifurcation point. In such cases, we slice the data about the point which is behind the tipping by approximately 20% of the respective pre-transition time series. On choosing a parameter (other than the control parameter) from a range of values, the bifurcation point varies and we have considered time series prior to the tipping, by taking into consideration the nearest tipping.

The negative noise autocorrelation (blue noise) results in the cancellation of short successive favourable and unfavourable conditions which doesn't have much effect on the tipping point. On the other hand, positive noise autocorrelation (red noise) leads to long runs of such events which bring forth a large amount of variability in systems and delay recovery (Yang et al 2019, van der Bolt et al 2018).

The EWSNet is trained using time series prior to the tipping whose endpoints precede the tipping by time units at least 20% of the original sequence length. For Dataset-C, the model is trained with time series generated from systems perturbed with correlated noise having correlation time $\kappa \in [-0.2, 0.2]$. The classification of transition variants for Dataset-C is a more difficult task compared to that of Dataset-W. Initially, we tried building the EWSNet with three convolution blocks, and 8 and 64 LSTM units, respectively. The performance of the EWSNet with 8 LSTM units (Figs. S1(a)-(c) and (g)-(i)) and 64 LSTM units (Fig. S1(d)-(f) and (j)-(l)) are presented suggesting the need for further hyper-parameter tuning. Confusion matrices show that EWSNet with 8 and 64 LSTM units result in a higher percentage of false alarms. The re-

| **Models** | **Parameters** | **Values** |
|---|---|---|
| [1] Saddle-node bifurcation: | $K$ - carrying capacity | 8–10 |
| $\frac{dN}{dt} = rN(1-\frac{N}{K}) - \frac{cN^2}{b^2+N^2}$ | $r$ - maximum growth rate | 1 |
| | $c$ -maximum grazing rate | **1–3** |
| | $b$ - half saturation constant | 1 |
| [2] Saddle-node bifurcation: | $K$ - carrying capacity | 11 |
| $\frac{dN}{dt} = N(1-\frac{N}{K}) - \frac{cN}{1+N}$ | $c$ - maximum grazing rate | **1–5** |
| [3] Saddle-node bifurcation: | $b$ - adult reproduction rate | 1 |
| $\frac{dJ}{dt} = bA - \frac{J}{1+J^2} - \mu_J J$ | $c$ - conversion rate | 1 |
| $\frac{dA}{dt} = \frac{J}{1+J^2} - AP - \mu_A A$ | $\mu_J$ - juvenile mortality rate | 0.05–0.1 |
| $\frac{dP}{dt} = APc - \mu_P$ | $\mu_A$ - adult mortality rate | 0.01 |
| | $\mu_P$ - predator mortality rate | **0.35–0.65** |
| [4] Saddle-node bifurcation: | $r$ - basal expression rate | 0.1 |
| $\frac{dx}{dt} = r + \frac{ax^2}{1+x^2} - x$ | $a$ - maximum transcription rate | **0–3** |
| [5] Saddle-node bifurcation: | $\mu$ - relative intensity of solar radiation | **0.5–2** |
| $\frac{dT}{dt} = \frac{1}{c}(-\epsilon\sigma T^4 + \frac{\mu I_o bT}{4} + \frac{\mu I_o(1-a)}{4})$ | $a$ -linear feedback | 2.81 |
| | $b$ -linear feedback | 0.009 |
| | $c$ - constant thermal inertia | $1.5\times10^8$ |
| | $\epsilon$ - effective emissivity | 0.69 |
| | $I_o$ - solar irradiance | 0.03 |
| [6] Transcritical bifurcation: | $K$ - carrying capacity | 10–15 |
| $\frac{dN}{dt} = rN(1-\frac{N}{K}) - cN$ | $r$ - maximum growth rate | 1 |
| | $c$ - maximum grazing rate | **0–2** |
| [7] Pitchfork bifurcation: | $K$ -carrying capacity | 8–10 |
| $\frac{dN}{dt} = rN(1-\frac{N}{K})(N-N_c) - cN + I$ | $r$ -maximum growth rate | **0.1–1** |
| | $c$ -maximum grazing rate | 0.8 |
| | $N_c$ - Allee threshold | 5 |
| | $I$ - immigration rate | 4 |
| [8] Supercritical Hopf bifurcation: | $K$ -carrying capacity of resource | **0.1–4** |
| $\frac{dN}{dt} = rN(1-\frac{N}{K}) - \frac{aNP}{b+N}$ | $r$ -maximum growth rate | 0.5 |
| $\frac{dP}{dt} = \frac{e_1 aNP}{b+N} - d_1 P$ | $a$ -maximum grazing rate | 0.4 |
| | $b$ - half saturation constant | 0.6 |
| | $e_1$ - assimilation efficiency of grazer | 0.6 |
| | $d_1$ - mortality rate of grazer | 0.15 |
| [9] No transition: | $K$ - carrying capacity | 1.9–3.9 |
| $\frac{dN}{dt} = rN(1-\frac{N}{K}) - \frac{cN^2}{b^2+N^2}$ | $r$ - maximum growth rate | 1 |
| | $c$ - maximum grazing rate | **1–3** |
| | $b$ -half saturation constant | 1 |

Table S1: **Model equations, parameters, and values:** Nine different deterministic base models and their parameter values were used to generate simulated time series for training and testing the EWSNet. For each model, the control parameter is shown in boldface with a range of values. To move a system across a bifurcation point (except model [9]), we gradually varied the control parameter across this range. In models [1], [3], [6], [7], and [9], apart from the control parameter, we choose another parameter from a given range of values to add more variations in the time series.

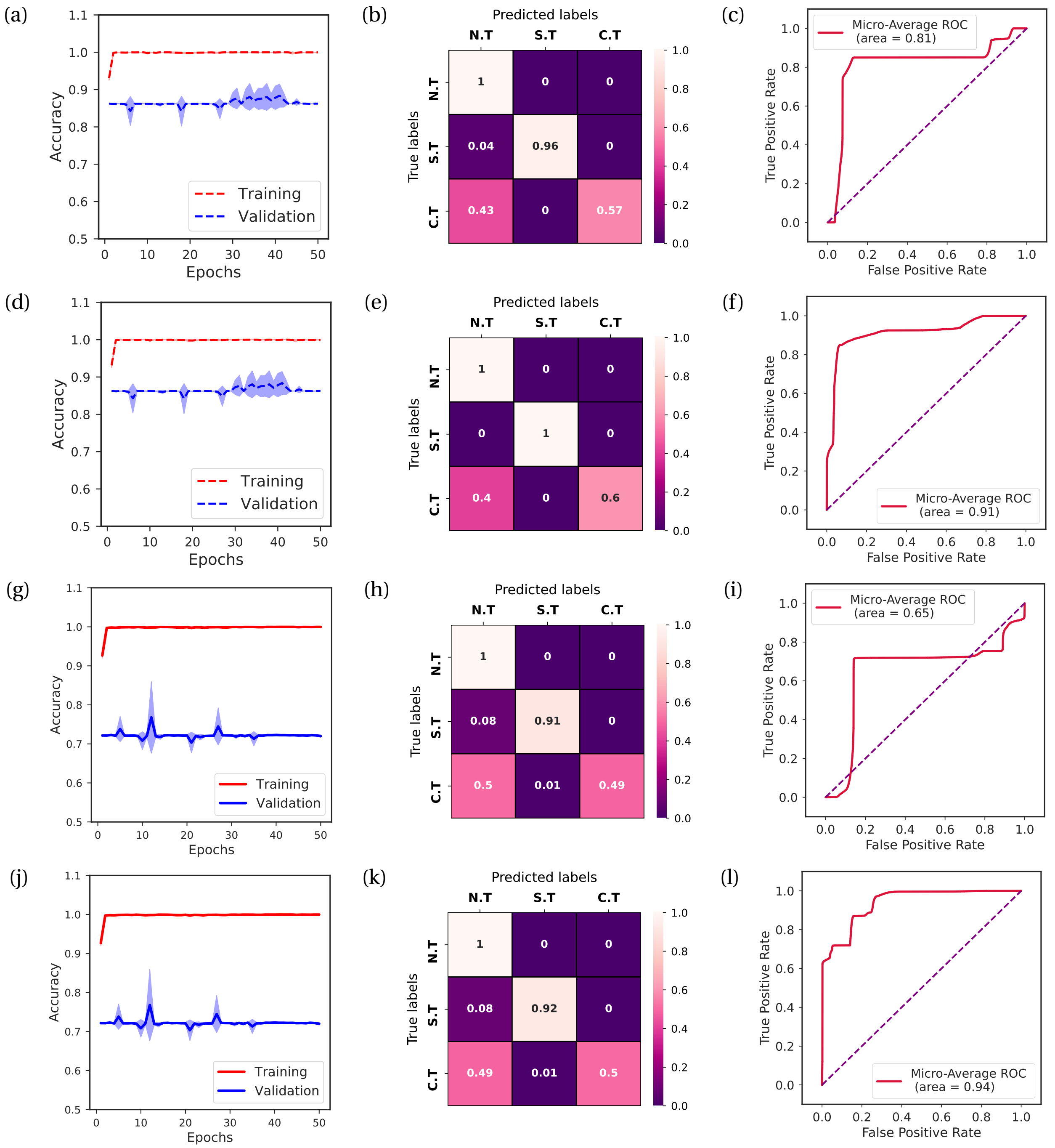


Figure S1: **Performance of the EWSNet on varying the model complexity:** (a,d) Accuracy, (b,e) confusion matrix, and (c,f) ROCs for the EWSNet corresponding to Dataset-W with (a-c) 8 LSTM units and (d-f) 64 LSTM units. Similarly, (g,j) accuracy, (h,k) confusion matrix, and (i,l) ROCs for the EWSNet corresponding to Dataset-C with (g-i) 8 LSTM units and (j-l) 64 LSTM units. Gradual improvement in the performance of the EWSNet on increasing model complexity (number of LSTM units) as observed from the confusion matrices and area under the ROCs indicate the need for a more complex model.

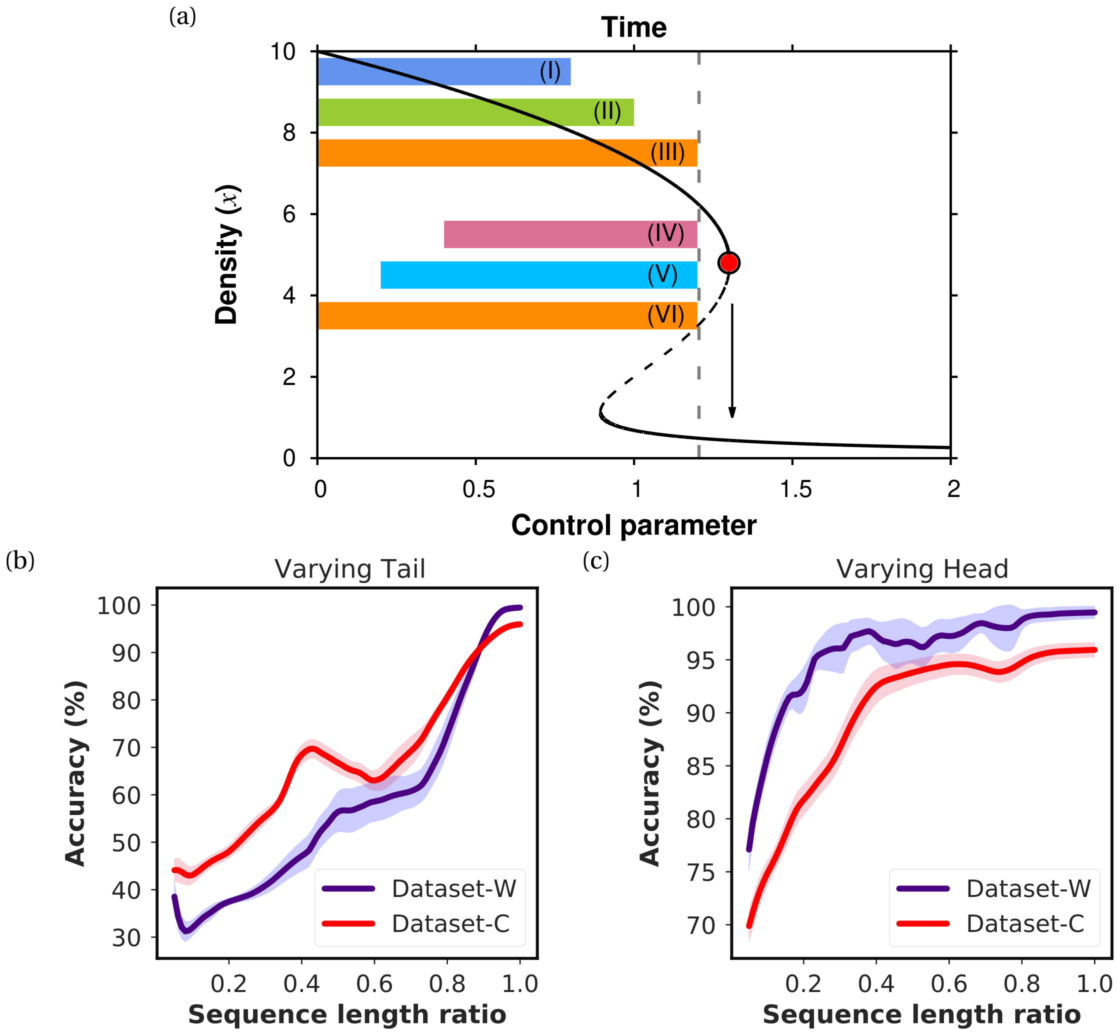


Figure S2: **Effects of varying tail and head of the time series on the final outputs of the EWSNet**: (a) A schematic figure representing two data resampling techniques used to study the efficacy of EWSNet; the bars with labels (I)-(III) denote varying tail, and labels (IV)-(VI) denote varying heads. The vertical dashed line represents the proximity to tipping up to which data is considered for training the EWSNet. The solid lines represent a stable equilibrium point and the dashed line represents an unstable equilibrium point. (b) Curves here show that the performance of EWSNet improves for time series with larger length, and which includes points closer to the tipping point. (c) Curves in this figure depict that keeping the distance from the tipping point fixed albeit increasing the length of the time series improves the performance of the EWSNet. Reducing the distance from tipping and increasing the time series length improves the performance of EWSNet.

ceiver operator curve (ROC) is a trade-off between the true positive and the false positive rates and the area under the ROC curve is representative of the model's efficiency in discriminating the classes. A model with a higher area under the ROC curve (close to 1) is a better classifier. Results indicate that a deep learning model with more hidden layers is required for the classification task which led us to model construction with three convolution blocks and 128 LSTM units (see main draft, Fig. 2).

To consider the effect of imperfect data sampling on the robustness of EWSNet we have resampled the time series with reduced time steps by randomly selecting out of the original ones followed by interpolation to the actual sequence length. This is done in order to eliminate the occurrence of artificial autocorrelation (Boettiger and Hastings 2012). Sampling is done in increment of 0.1 starting from 10% to full length. The resampled time series are further passed to the pretrained EWSNet to test its efficiency concerning imperfectly sampled data. We have also studied the effect of missing data closer and away from the tipping on the EWSNet in classifying the transitions. To account for this, we have sliced data points just prior to tipping (varying tail) or initial dynamics (varying head) of different lengths and computed the accuracy of the EWSNet (see Fig. S2). Results indicate the improved performance of EWSNet as we incorporate time series with endpoints closer to the tipping. Additionally, increasing the length of the time series improves the accuracy of the EWSNet. The EWSNet is implemented using the Tensorflow framework (Abadi et al 2015).

### S2. Data generation for classical ML models trained on generic EWSs

To study whether classical ML models (see Table S2) trained with generic EWSs can classify a transition or not, we compute generic EWSs for combinations of bandwidths and window sizes (for details see "Methods: Machine learning models trained on generic EWSs" in the main draft). They are further concatenated for each generic EWS, e.g., standard deviation (SD), autocorrelation at lag-1 (AR-1), skewness, kurtosis, auto-regressive coefficient at lag-1 (ACF-1), return rate (RR), density ratio (DR), coefficient of variation (CV) and their pairwise combinations, and treated as an attribute based on which the classical ML models infer the type of a transition ahead in a sequence of events.

Results of early warning indicators are majorly dependent on the choice of window size and filtering bandwidth. A choice of short sliding window leads to undulating indicator trends whereas longer ones result in flatter estimates. Considering narrower filtering bandwidth, get rid of low frequencies, which are essential in capturing the CSD trends. Whereas a choice of broader filtering bandwidth fails to remove the invariabilities which may cause factitious autocorrelation. Hence, we choose a range of bandwidths and window sizes to calculate generic EWSs which are further used to train the classical ML models for improved performance (Lenton et al 2012, Boettiger and Hastings 2012). The results for combined generic EWSs passed as input to the classical ML models are presented in (Figs. S3-S4). Among all the generic EWSs and their combinations, AR1+SD anticipates transitions with minimal false alarms (Fig. S5).

| **ML models** | **Principle** | **Parameters** |
|---|---|---|
| Logistic Regression (LR) | For a given set of features, the model performs the classification task by estimating probabilities using the logistic function post linear regression fitting. | regularizer (C) = 0.5, penalty = cross entropy, solver = 'sag' |
| Random Forest (RF) | Ensemble learning algorithm that consist of a host of decision trees. A prediction is obtained from every decision tree and the mode of the resulting prediction distribution is considered as final output. | trees = 100, penalty = gini |
| Support Vector Machine (SVM) | A non-probabilistic approach that learns the maximum margin hyperplane fitting the empirical data distribution. It classifies the different classes after mapping the n-dimensional input features to points in a latent space using a kernel requisite to the task as per the separability of the input space. | Kernel = linear, penalty = hinge loss, regularizer = 0.5 |
| Multilayer perceptron (MLP) | MLP is a class of feed-forward artificial neural networks composed of multiple layers of perceptrons. It learns a function $f : R^m \rightarrow R^o$ by training on an m dimensional input space and o is the dimension of the output space. Uses nonlinear activation function and backpropagation to efficiently learn weights corresponding to classes in a classification task. | penalty = cross entropy loss, optimizer = Adam, hidden layers = 1, No. of nodes per hidden layer = 100 |

Table S2: **Classical ML models used for characterizing trends in generic EWSs**: Brief description of classical ML models and the parameter values for the respective. All the classical ML models are implemented using Scikit-learn package (Pedregosa et al 2011).

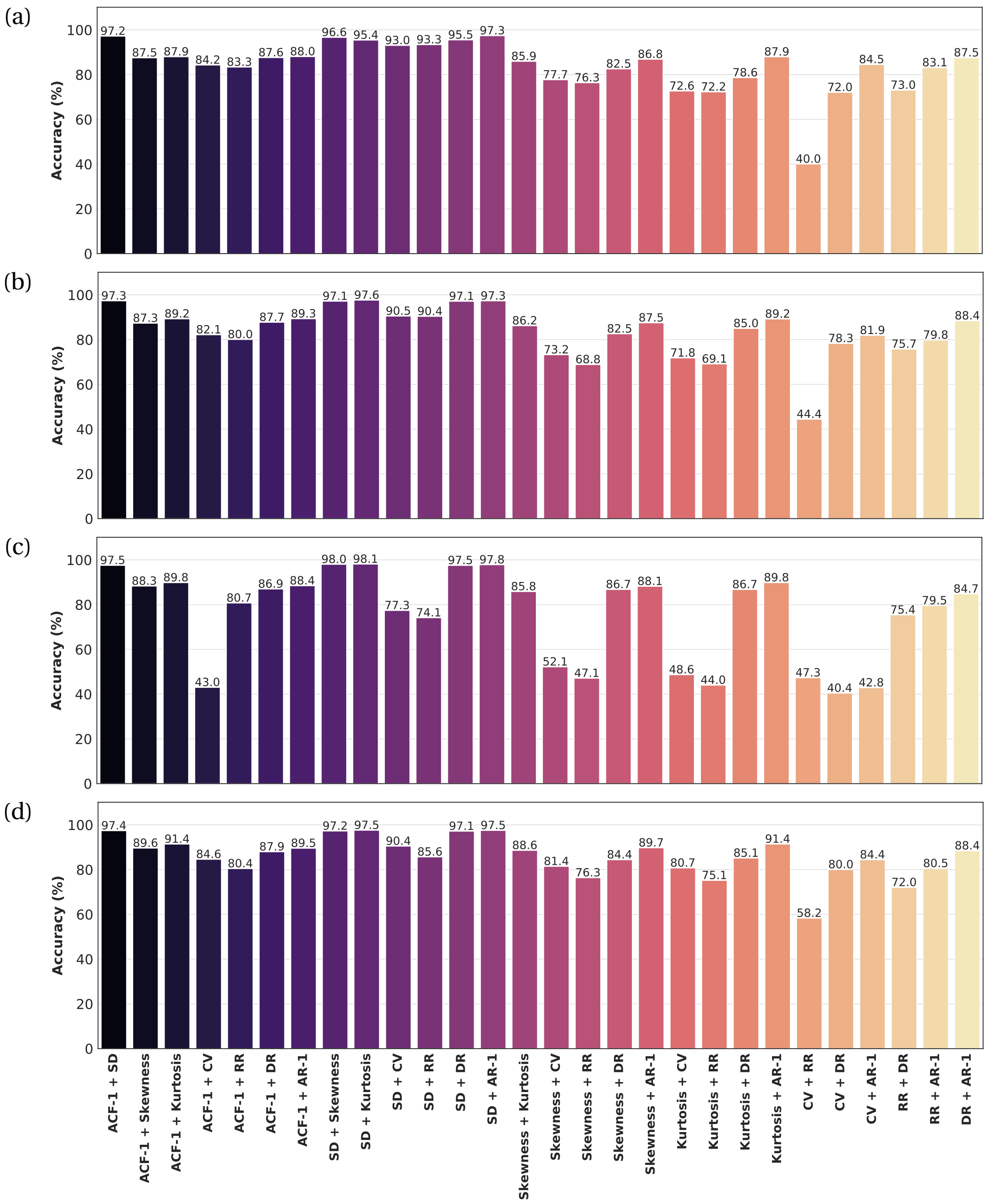


Figure S3: **Performance of generic EWSs using four standard ML models for Dataset-W:** (a) Logistic Regression, (b) Support Vector Machine, (c) Random Forest, and (d) Multilayer perceptron. Combined EWSs for a combination of window sizes and bandwidths are used to train the ML models and the output results as bar plots corresponding to each indicator. SD+AR1 performs best among all the $\binom{8}{2}$ combinations of the 8 leading generic indicators irrespective of the chosen classical ML model.

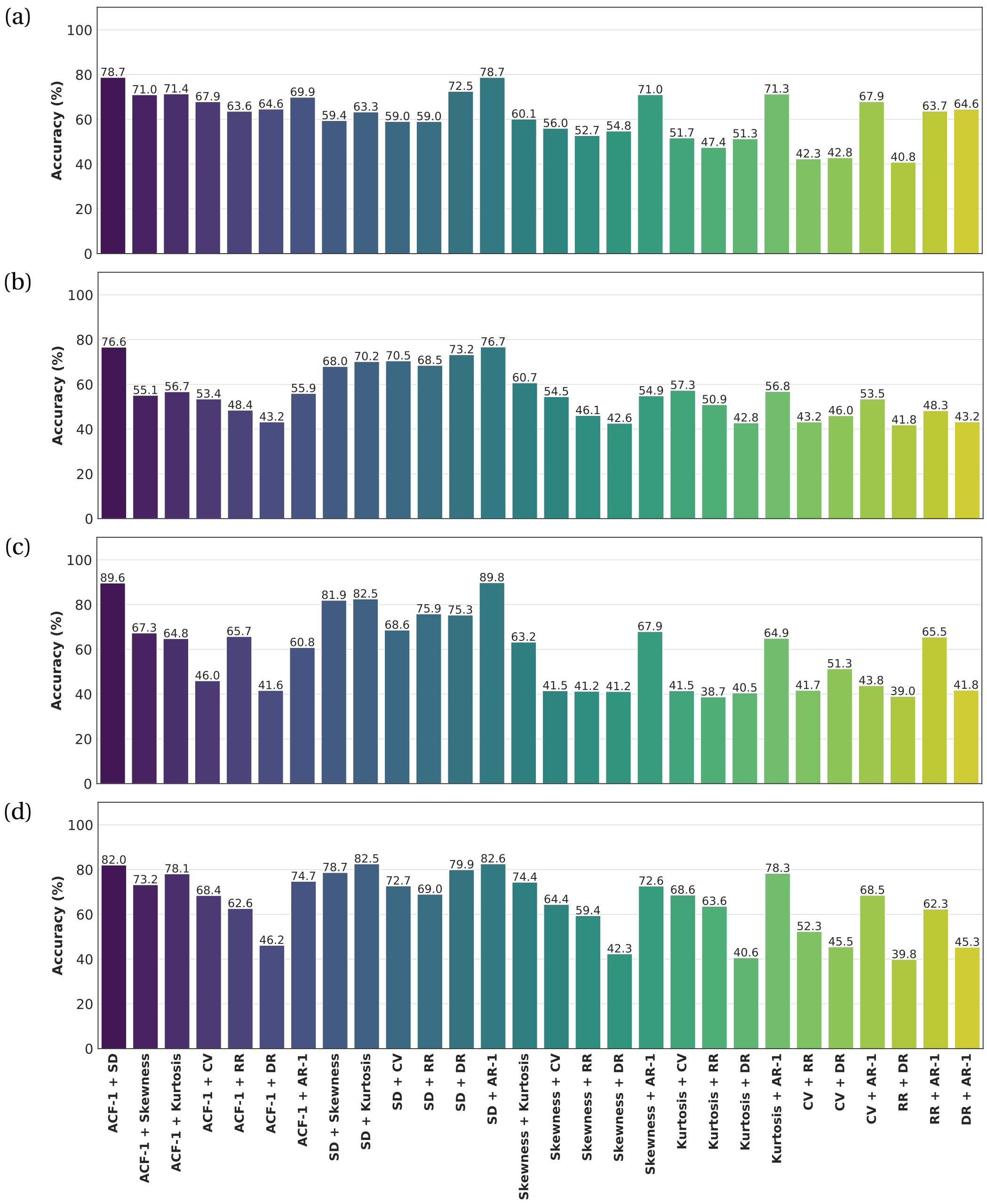


Figure S4: **Performance of generic EWSs using four standard ML models for Dataset-C:** (a) Logistic Regression, (b) Support Vector Machine, (c) Random Forest, and (d) Multilayer perceptron. Combined EWSs for a combination of window sizes and bandwidths are used to train the ML models and the output results as bar plots corresponding to each indicator. SD+AR1 performs best among all the $\binom{8}{2}$ combinations of the 8 leading generic indicators irrespective of the chosen classical ML model.

### S3. Real data description

To analyze the practicability of EWSNet in anticipating real world transitions, we validated our model on the seventeen available real data of catastrophic and non-catastrophic shifts (8

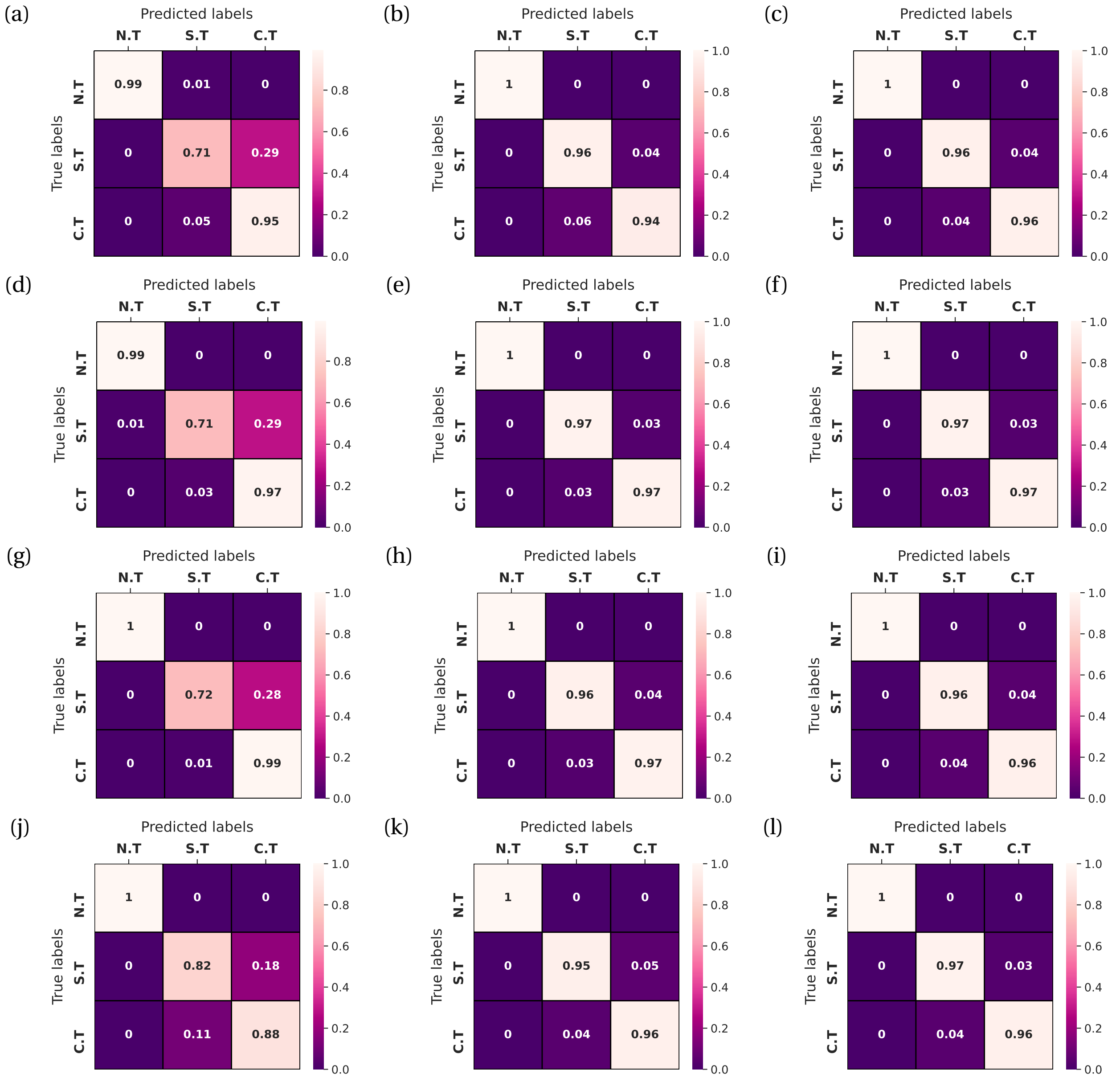


Figure S5: **Confusion matrices for classical ML models to study the effectiveness of generic EWSs**: Confusion matrices corresponding to autocorrelation at lag-1, standard deviation and their combination, for (a-c) Logistic Regression, (d-f), Random Forest, (g-i) Support Vector Machine, and (j-l) Multilayer perceptron, respectively.

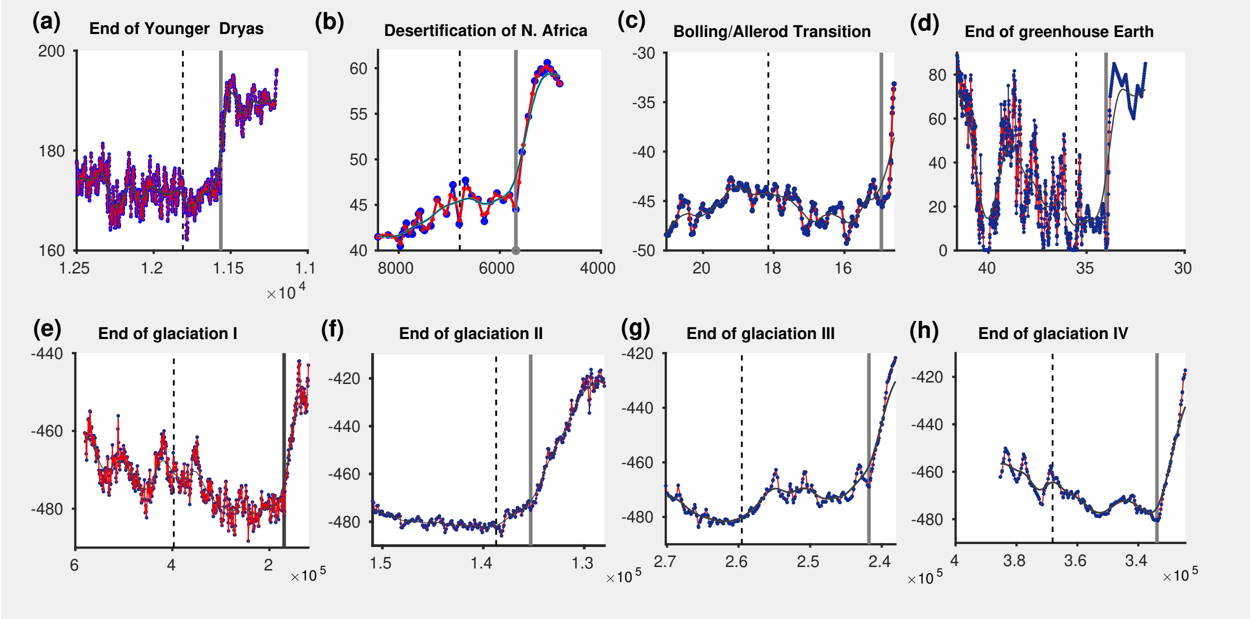


Figure S6: **Real paleoclimatic data exhibiting sudden climatic transitions** : Reconstructed time series of abrupt climatic shifts. The vertical dashed line in each sub-figure indicates the time series considered for calculating EWSs, the grey band marks the transition region, and the black curve through each time series represents the Gaussian kernel fitted to filter out trends.

paleoclimatic and 9 bio-ecological time series). The paleoclimatic data includes a range of abrupt climate shifts that correspond to diverse real climate systems (Fig. S6). These transitions include the end of Younger Dryas (Hughen et al 2000), desertification of North Africa (DeMenocal 2001), Bolling Allerod transition (Alley et al 2004), end of greenhouse earth (Tripati et al 2005), end of Glaciation 1, 2, 3, 4 (Vimeux et al 2001). The end of the Younger Dryas about 11,500 years ago was the return to the glacial phase after the interstadial resulting in a decline in the temperature of Greenland by $4-10^{o}$C and prevailing drier conditions. The Bolling Allerod transition is the abrupt warm and moist interstadial period that occurred during the last glaciation period studied using the 100,000 years old oxygen isotope. End of Greenhouse earth represents the transition from extreme global warmth (55 million years ago) to the present glaciated state. These are some of the most prominent abrupt climatic shifts and have therefore been used to study the efficacy of the EWSNet.

The other time series include population collapse in deteriorating environment (Clements and Ozgul 2016), collapse of whale stock over the 20th century (Clements et al 2017), and

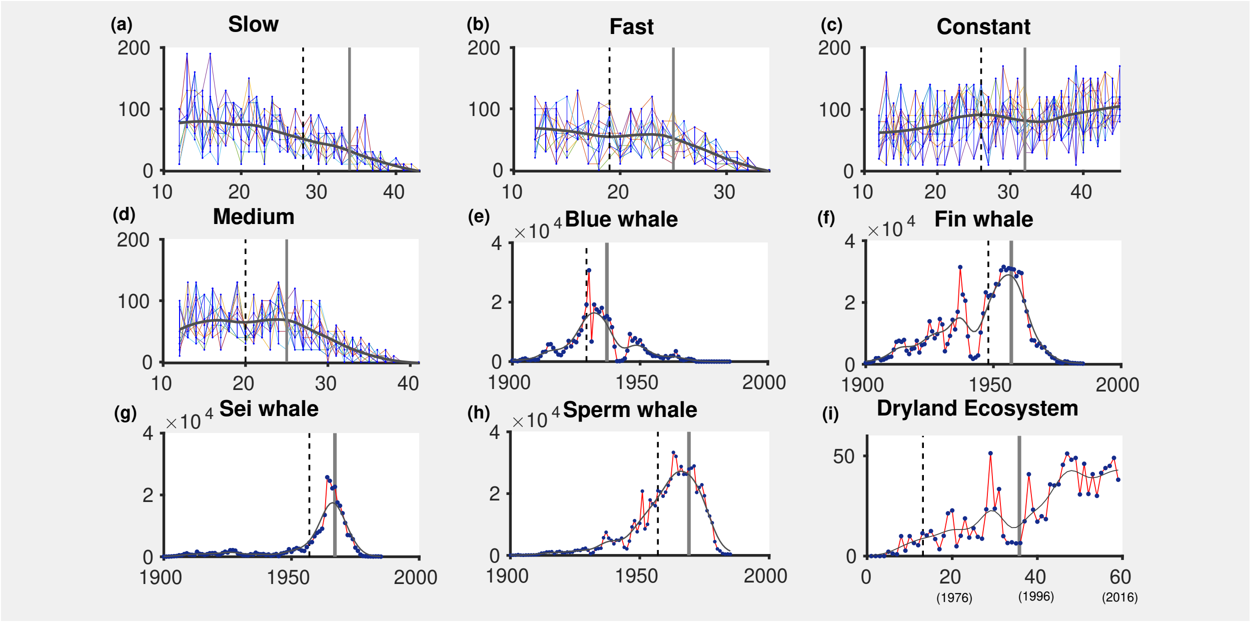


Figure S7: **Ecological and experimental data depicting transitions** : (a-d) Data on *Didinium nasutum* mean body size exposed to 4 different mediums; viz slow, fast, constant, and medium. Black curve through time series represents LOESS (locally weighted smoothing) smoothing showing the average change in mean body size of *Didinium nasutum*. (e-h) Time series depicting the change in whale count caught over the given period of time. The black curve represents LOESS smoothing to visualize mean trends in whale abundance. (i) Catastrophic transition in the vegetative state of a dryland ecosystem, the black line indicates Gaussian kernel to filter out slower trends.

abrupt shifts in a dryland ecosystem (Chen et al 2018) (Fig. S7). All the time series have been interpolated to fill missing values and consideration of equidistant time steps. This is done in accordance with the original time series allowing no contradictory trends due to interpolation. This is followed by Gaussian filtering for the paleoclimatic and dryland ecosystem data and LOESS smoothing for the rest to view the overall trend in the time series. Further computing the AR-1 taking window size half of the length of the time series and null bandwidth to compare the performance and robustness of the traditional and modern ML approach. Table S3 presents Kendall's $\tau$ values for generic EWSs and prediction probability for the EWSNet.

| **Reconstructed time series of various transitions** | **EWS signal** | **Kendall's-$\tau$** | **EWSNet** | **Prediction probability** |
|---|---|---|---|---|
| End of Younger Dryas (Hughen et al 2000) | strong | 0.69 | incorrectly classified | 0.59 |
| Desertification of North Africa (De-Menocal 2001) | moderate | 0.59 | incorrectly classified | 0.53 |
| Bolling Alleroid transition (Alley et al 2004) | weak | 0.27 | correctly classified | 1 |
| End of Greenhouse Earth (Tripati et al 2005) | weak | 0.83 | correctly classified | 0.89 |
| Glaciation 1 (Vimeux et al 2001) | strong | 0.8 | correctly classified | 1 |
| Glaciation 2 (Vimeux et al 2001) | weak | 0.17 | correctly classified | 1 |
| Glaciation 3 (Vimeux et al 2001) | moderate | 0.43 | correctly classified | 1 |
| Glaciation 4 (Vimeux et al 2001) | moderate | 0.52 | correctly classified | 1 |
| *D. nasutum* in slow (Clements and Ozgul 2016) | weak | 0.143 | correctly classified | 0.43 |
| *D. nasutum* in fast (Clements and Ozgul 2016) | weak | 0.143 | incorrectly classified | 0.83 |
| *D. nasutum* in constant (Clements and Ozgul 2016) | weak | 0.06 | correctly classified | 0.97 |
| *D. nasutum* in medium (Clements and Ozgul 2016) | weak | 0.048 | incorrectly classified | 0.92 |
| Blue whale (Clements et al 2017) | weak | 0.226 | correctly classified | 1 |
| Fin whale (Clements et al 2017) | strong | 0.688 | correctly classified | 1 |
| Sei whale (Clements et al 2017) | weak | 0.162 | correctly classified | 1 |
| Sperm whale (Clements et al 2017) | weak | 0.313 | correctly classified | 1 |
| Dryland Ecosystem (Chen et al 2018) | strong | 0.867 | correctly classified | 0.99 |

Table S3: **Robustness of AR-1 and EWSNet in forewarning transitions in real and experimental data sets**: Classification of real time series using the EWS AR-1 and the EWSNet. Reliability of the signals are measured using Kendall's-$\tau$ for AR-1 and prediction probability for EWSNet.
